# Interspecies variation of 45S ribosomal DNA in vertebrates

**DOI:** 10.64898/2026.06.09.731232

**Authors:** Lin Kang, Giulio Formenti, Treyton O’Connor, Pawel Michalak, the Vertebrate Genomes Project Consortium Phase I

**Affiliations:** Edward Via College of Osteopathic Medicine, Monroe, LA 71203, USA; Center for One Health Research, VA-MD Regional College of Veterinary Medicine, Blacksburg, VA 24060, USA; College of Pharmacy, University of Louisiana Monroe, Monroe, LA 71203, USA; The Vertebrate Genome Laboratory, The Rockefeller University, New York, NY 10065, USA; Institute of Evolution, University of Haifa, Haifa, Israel

## Abstract

Ribosomal DNA (rDNA) remains one of the least resolved components of complex genomes despite its central role in cellular function and evolutionary inference. Using chromosome-scale assemblies from the Vertebrate Genomes Project, we present a comparative analysis of the 45S rDNA locus across vertebrates. rDNA organization varies widely among clades, with lineage-specific differences in chromosomal distribution, copy number, and positional bias. Mammals and birds typically restrict rDNA to few, often terminal loci, whereas amphibians and several fish lineages exhibit highly dispersed architectures, including extreme cases spanning dozens of chromosomes. Copy number varies by more than an order of magnitude and does not scale simply with repeat-unit size or predicted functional demand, consistent with rDNA functioning as a flexible genomic reservoir rather than a direct proxy for ribosome production. At the sequence level, the 45S unit resolves into distinct evolutionary regimes. Genes encoding rRNAs (18S, 5.8S, 28S) remain highly conserved and retain strong phylogenetic signal, whereas internal transcribed spacers show rapid divergence, lineage-specific expansion, and pronounced asymmetry between intergenic spacers ITS1 and ITS2. Sequence–structure analyses reveal a gradient of constraint across the locus, from tightly coupled evolution in rRNA-encoding regions to increasingly permissive regimes in spacers. Structure-informed phylogenetic inference yields modest but consistent improvements under high divergence, highlighting its value when sequence signal degrades. Together, these results establish the 45S rDNA locus as a multi-layered evolutionary system integrating deep conservation with structural plasticity across vertebrates.

## Introduction

Ribosomal RNA genes occupy a uniquely central position in both cellular metabolism and evolutionary biology. As the structural and catalytic core of the ribosome, rRNAs are indispensable to protein synthesis and represent the most abundant transcripts in all living cells, accounting for up to 80–90% of total RNA in vertebrate transcriptomes^1,2^. Their evolutionary origins extend to the last universal common ancestor (LUCA), placing rRNA among the most ancient and deeply conserved genetic sequences known^3^. This extraordinary conservation, particularly of the 18S, 5.8S, and 28S rRNAs, has made ribosomal genes foundational markers for reconstructing deep phylogenetic relationships across all domains of life, including the major lineages of vertebrates^4^.

In eukaryotes, rRNA genes are organized into large tandem arrays of 45S rDNA, typically comprising hundreds to thousands of repeat units that form the nucleolar organizer regions (NORs). Each repeat unit encodes the 18S, 5.8S, and 28S rRNAs, separated by two internal transcribed spacers, ITS1 and ITS2, which are transcribed as part of a single precursor and subsequently excised during rRNA maturation (**Fig. 1**). Despite their centrality to genome function, rDNA arrays have long remained among the least characterized regions of complex genomes. Their extreme repetitiveness, multilocus organization, high GC content, and dynamic copy-number variation have historically rendered them refractory to assembly and analysis, leading to their systematic exclusion or collapse in reference genomes and their characterization as genomic “dark matter”^5^.

**Figure 1.**
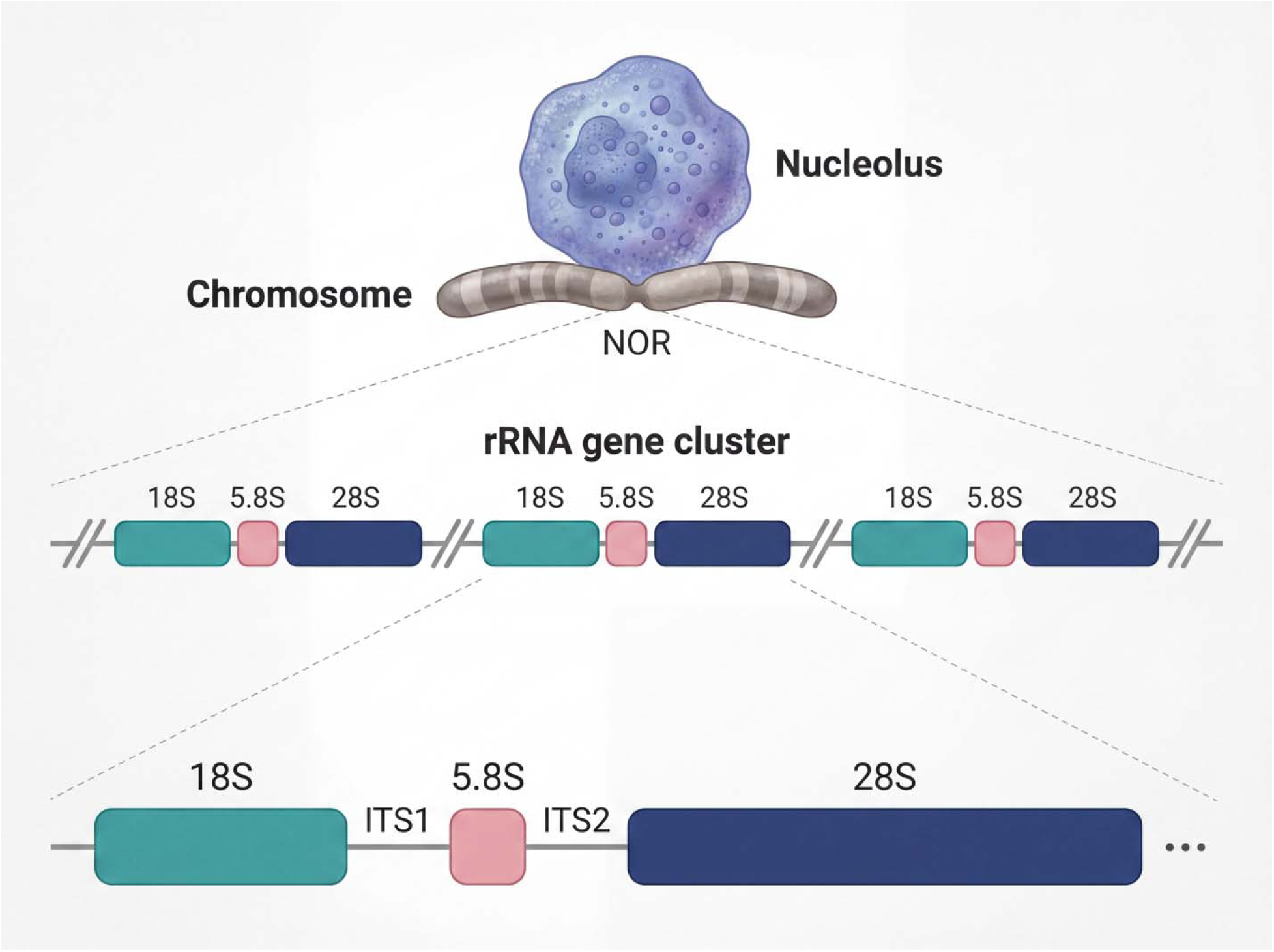
Genomic organization of the vertebrate 45S rDNA locus. Schematic illustrating the hierarchical organization of ribosomal DNA within the nucleus. Ribosomal RNA genes are localized to the nucleolus and encoded within nucleolar organizer regions (NORs) on chromosomes, where they occur as tandemly repeated arrays of 45S rDNA units. Each repeat comprises the 18S, 5.8S and 28S rRNA encoding genes separated by the internal transcribed spacers ITS1 and ITS2, arranged in the canonical order 18S–ITS1–5.8S–ITS2–28S. The diagram depicts the multi-scale architecture from chromosomal localization to the structure of a single transcriptional unit.

In humans, the 45S rDNA clusters are located on the short (p) arms of the five acrocentric chromosome pairs: 13, 14, 15, 21, and 22. Even in the first complete assembly of a human genome from the Telomere-to-Telomere (T2T) Consortium, which resolved centromeres, acrocentric short arms, and other previously inaccessible repeats^6^, the rDNA sequences were only represented as a model. Overall, accurate characterization of rDNA copy number, sequence variation, and evolutionary dynamics remains an active area of investigation, highlighting rDNA as one of the final frontiers in genome completeness^6–9^.

Beyond their role as ribosome templates, rRNA genes and the nucleolus form a highly organized and functionally specialized regulatory system. Nucleolar assembly is driven by rDNA transcription itself, and rDNA is regulated by a dedicated RNA polymerase I machinery, specialized chromatin states, and distinct replication and epigenetic dynamics that distinguish it from most other genomic loci^10–12^. Although only a subset of rRNA gene copies is transcriptionally active at any given time, rDNA arrays exhibit remarkable structural persistence, concerted evolution, and epigenetic memory across cell divisions^8,13,14^. Experimental reductions in rDNA copy number often have surprisingly modest effects on organismal viability and bulk protein synthesis^15^, suggesting that rDNA redundancy exceeds minimal translational requirements and may instead provide regulatory buffering or evolutionary flexibility, including the phenomenon of nucleolar dominance during hybridization^16,17^.

Within this uniquely constrained yet dynamic genomic environment, the evolutionary behavior of internal transcribed spacers is striking. In contrast to the near invariance of rRNA coding sequences, ITS regions evolve rapidly, showing extensive length variation, dramatic shifts in nucleotide composition, and little primary-sequence conservation even among closely related species. This pattern is generally attributed to concerted evolution, whereby unequal crossing-over and gene conversion homogenize rDNA copies within species while allowing rapid divergence between lineages^18,19^. As a result, ITS sequences have become widely used as high-resolution molecular markers for species identification and shallow phylogenetic inference across eukaryotes^20–22^.

Despite extensive use of ITS regions in systematics, their large-scale evolutionary properties across vertebrates remain poorly characterized. Previous studies have typically focused on limited taxonomic sampling, individual clades, or single ITS regions, leaving unresolved how spacer length, nucleotide composition, and divergence scale across the vertebrate tree. Moreover, ITS1 and ITS2 are often implicitly treated as a single evolutionary unit, despite evidence that they differ in length distributions, compositional biases, secondary structure constraints, and evolutionary trajectories^23,24^. Addressing these gaps requires both dense phylogenetic sampling and genome assemblies capable of resolving full-length rDNA repeats, conditions that have only recently been met. Here, we exploit chromosome-scale assemblies generated by the Vertebrate Genomes Project (VGP), which aims to produce error-corrected, haplotype-resolved reference genomes across all extant vertebrate orders^25^. Using a uniform annotation pipeline and stringent quality filtering, we systematically identify and characterize 45S rDNA transcription units, along with ITS1 and ITS2 sequences across vertebrates and describe their unique evolutionary patterns.

## Results

### Chromosomal distribution of rDNA arrays differs sharply among vertebrate clades

The number of chromosomes harboring 45S rDNA loci varies substantially across clades (**Fig 2a; Supplementary Tables S1 & S2**). In most mammals, birds, lepidosaurs, turtles, and many fishes, rDNA clusters are confined to one or two chromosomes. In contrast, amphibians show a median of approximately five chromosome-level molecules carrying rDNA loci, and the sampled lobe-finned fishes show even broader dispersion (**Fig 2b**). The lungfish *Protopterus annectens* represents an extreme case, with 45S rDNA detected on 26 chromosome-level molecules, indicating a highly dispersed rDNA architecture.

**Figure 2.**
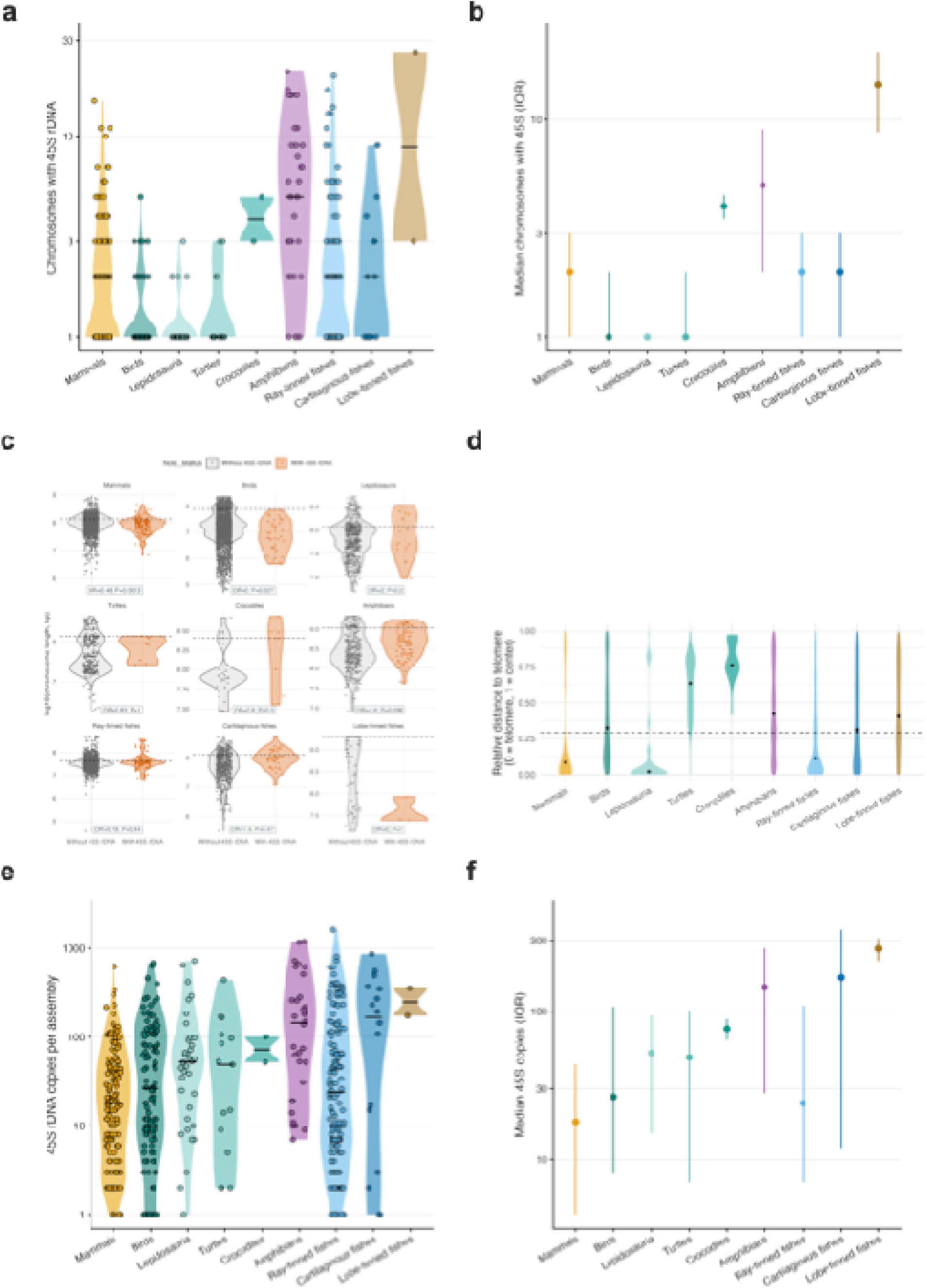
Chromosomal distribution across vertebrate clades and variation in 45S rDNA copy number. (**a**) Number of chromosomes harboring 45S rDNA loci per assembly. Most taxa possess rDNA on one or a few chromosomes, but some clades—notably amphibians and lobe-finned fishes—show expansion to multiple chromosomes. (**b**) Median number of 45S-bearing chromosomes per clade with IQR, highlighting lineage-specific differences in chromosomal distribution, from predominantly single-locus organization in amniotes to more dispersed arrangements in amphibians and fishes. **(c)** Comparison of chromosome lengths for chromosomes with (orange) versus without (grey) 45S rDNA, stratified by clade. For each clade, points represent individual chromosomes and violins summarize the length distribution. The dashed horizontal line marks the clade-specific N50 chromosome length, defined as the length above which 50% of the total chromosomal sequence is contained. Text within each facet reports the Fisher’s exact odds ratio (OR) and P value testing enrichment of 45S-bearing chromosomes among “large” chromosomes (length ≥ N50) versus “small” chromosomes (length < N50). Across clades, 45S rDNA loci are non-randomly distributed with respect to chromosome size, frequently occurring on chromosomes near or above the N50, although the strength and direction of this bias vary among lineages. **(d)** Relative telomeric positioning of 45S rDNA arrays by clade. For each 45S array, the position along its host chromosome was converted to a scaled distance from the nearest telomere (tel_frac; 0 = at the telomere, 1 = at the chromosome midpoint). Violin plots show the distribution of tel_frac values within each clade. The dashed line marks the expected median tel_frac under a null model of uniformly random positions along a chromosome (∼0.29). Across clades, 45S rDNA loci are frequently enriched toward chromosomal ends, with strong telomeric bias in mammals, lepidosaurs and ray-finned fishes, where distributions are skewed toward low values. In contrast, turtles, amphibians and lobe-finned fishes show broader or more centrally distributed patterns, indicating reduced or heterogeneous positional bias. The terminal region (tel_frac ≤ 0.1) defines the threshold used for enrichment testing. These results indicate that while telomeric localization of 45S rDNA is common across vertebrates, the strength of this bias varies markedly among lineages. (**e**) Distribution of 45S rDNA copy number per genome assembly across major vertebrate clades. Points represent individual assemblies plotted on a logarithmic scale, with violins indicating density. Copy number varies over several orders of magnitude within and among clades, with particularly broad dispersion in ray-finned fishes and amphibians. (**f**) Median 45S rDNA copy number per clade with interquartile range (IQR). Central tendencies differ substantially among lineages, with higher median values in lobe-finned fishes, cartilaginous fishes and amphibians compared to mammals and birds.

Testing whether rDNA clusters preferentially occur on large chromosomes revealed strong clade-specific differences rather than a universal rule (**Fig 2c**). In mammals, 45S loci are significantly depleted on large chromosomes, both in hosting frequency and in copy density. Birds show an even stronger depletion, with no large chromosomes carrying rDNA under this classification scheme. Similar depletion patterns are observed in ray-finned fishes and turtles, which exhibit lower copy density of rDNA on large chromosomes. In contrast, amphibians show a significant enrichment of both hosting frequency and repeat density on large chromosomes, whereas lepidosaurs exhibit strongly elevated copy density on large chromosomes despite only modest enrichment in host frequency.

These findings indicate that the chromosomal ecology of rDNA arrays varies substantially among vertebrate lineages and likely reflects lineage-specific differences in recombination dynamics, chromosomal rearrangements, and nucleolar organizer region evolution.

Analysis of the relative distance of 45S rDNA arrays to chromosome ends revealed a pronounced tendency for rDNA loci to occur in terminal regions in several vertebrate clades, although the strength of this pattern varies markedly across lineages (**Fig. 2d**). Mammals show the strongest telomeric bias, with most loci clustering very close to chromosome ends and a large proportion falling within the defined terminal interval (tel_frac ≤ 0.1), producing a clear enrichment relative to the random expectation. Ray-finned fishes and Lepidosauria also exhibit a noticeable terminal skew, whereas birds, turtles, and amphibians display broader distributions extending toward more internal chromosomal positions. In several non-tetrapod lineages, including cartilaginous and lobe-finned fishes, rDNA loci occur across a wide span of distances from the telomere, suggesting weaker positional constraints. The pronounced terminal enrichment in mammals is consistent with the well-known tendency of ribosomal DNA arrays to occupy the short arms of acrocentric chromosomes, particularly in primates, where nucleolar organizer regions are typically located on the p arms of chromosomes such as human 13, 14, 15, 21, and 22^26^. These short arms are themselves immediately adjacent to telomeres, effectively placing rRNA gene clusters in a terminal chromosomal context. Thus, while terminal localization of 45S rDNA is not universal across vertebrates, the mammalian, and especially primate, pattern supports the idea that structural features of acrocentric chromosomes strongly influence the genomic positioning of nucleolar organizer regions.

### Copy number of 45S rDNA repeats varies by more than an order of magnitude among vertebrates

The number of inferred 45S rDNA repeat pairs varies dramatically across vertebrate genomes, spanning more than an order of magnitude among the sampled taxa. Median copy numbers are highest in amphibians, cartilaginous fishes, and the two sampled lobe-finned fishes, whereas mammals, birds, and most ray-finned fishes generally harbor substantially fewer copies (**Fig. 2e-f; Supplementary Table S3**). In many mammalian and avian genomes, inferred repeat numbers fall within a relatively modest range of several tens of pairs, consistent with cytogenetic estimates of nucleolar organizer regions in these clades^27^. By contrast, amphibians frequently exhibit much larger arrays, reflecting the well-known tendency of their genomes to contain extensive repetitive DNA and large rDNA clusters. Several species represent striking outliers within the dataset. The teleost *Coregonus lavaretus* shows an estimated 1,625 paired 45S repeats, representing the largest inferred rDNA array among the vertebrates analyzed. Similarly elevated copy numbers occur in amphibians such as *Triturus cristatus* and *Lissotriton helveticus*, each exceeding 1,000 inferred repeat pairs. These extreme values contrast sharply with the smaller arrays observed in many mammals and birds.

Variation in rDNA copy number may also have functional implications for nucleolar activity and ribosome biogenesis capacity, as the number of rDNA repeats determines the potential number of transcriptionally active rRNA gene copies available to support ribosome production. However, rDNA arrays are typically only partially active, with a substantial fraction of repeats remaining epigenetically silenced^8,28^. Consequently, increases in copy number do not necessarily translate directly into increased rRNA transcription but may instead provide a buffer against genomic instability or recombination-mediated losses within the array^29^. It is important to interpret absolute repeat numbers with caution. Accurate estimation of rDNA copy number remains technically challenging, even in high-quality assemblies, as rDNA arrays consist of long, highly homogeneous tandem repeats that are difficult to assemble and quantify precisely. Although long-read sequencing and chromosome-scale assemblies such as those generated by the Vertebrate Genomes Project substantially improve representation of repetitive regions, repeat counts may still be influenced by assembly completeness, repeat collapsing, or unresolved array boundaries. Thus, while the broad patterns reported here are robust, the absolute copy numbers for individual species should be interpreted as approximate estimates rather than exact counts.

### rDNA repeat-unit size varies widely across vertebrates and is driven primarily by spacer architecture

Comparative analysis revealed substantial differences in the size of the full rDNA repeat unit across vertebrate clades (**Fig. 3a-f; Supplementary Table S3**). Median repeat-unit lengths were largest in mammals (∼34.5 kb) and somewhat smaller in birds (∼21 kb), turtles (∼20 kb), and cartilaginous fishes (∼20.8 kb). Shorter repeats were observed in amphibians (∼12.7 kb) and ray-finned fishes (∼14.1 kb). Importantly, rDNA repeat-unit size or spacer length do not scale directly with repeat copy numbers. Lineages with the largest ITS expansions, such as mammals and birds, often possess comparatively modest repeat numbers, whereas clades with compact spacers, including amphibians and several fishes, frequently maintain very large rDNA arrays. This decoupling indicates that repeat-unit architecture and array copy number represent largely independent axes of rDNA evolution, likely shaped by lineage-specific dynamics of unequal crossing-over, gene conversion, and the chromosomal stability of nucleolar organizer regions. Correlation analysis indicates that repeat-unit expansion is largely driven by variation in spacer length rather than changes in the rRNA coding regions. Total rDNA repeat length shows its strongest correlation with ITS2 length (Pearson r= 0.56) and a weaker correlation with ITS1 length (r= 0.23), whereas correlations with genome size are negligible or slightly negative (**Fig. 3g**). These results indicate that rDNA repeat expansion reflects local architectural changes within the rDNA locus rather than genome-wide scaling of repetitive DNA. Several species represent extreme repeat-length outliers. For example, deer *Dama dama* exhibits a median rDNA repeat length of ∼90.7 kb, while tropicbird *Phaethon aethereus* and teleost *Parambassis ranga* show repeat units exceeding 84.5 kb and 57.6 kb respectively (**Supplementary Table S3**).

**Figure 3.**
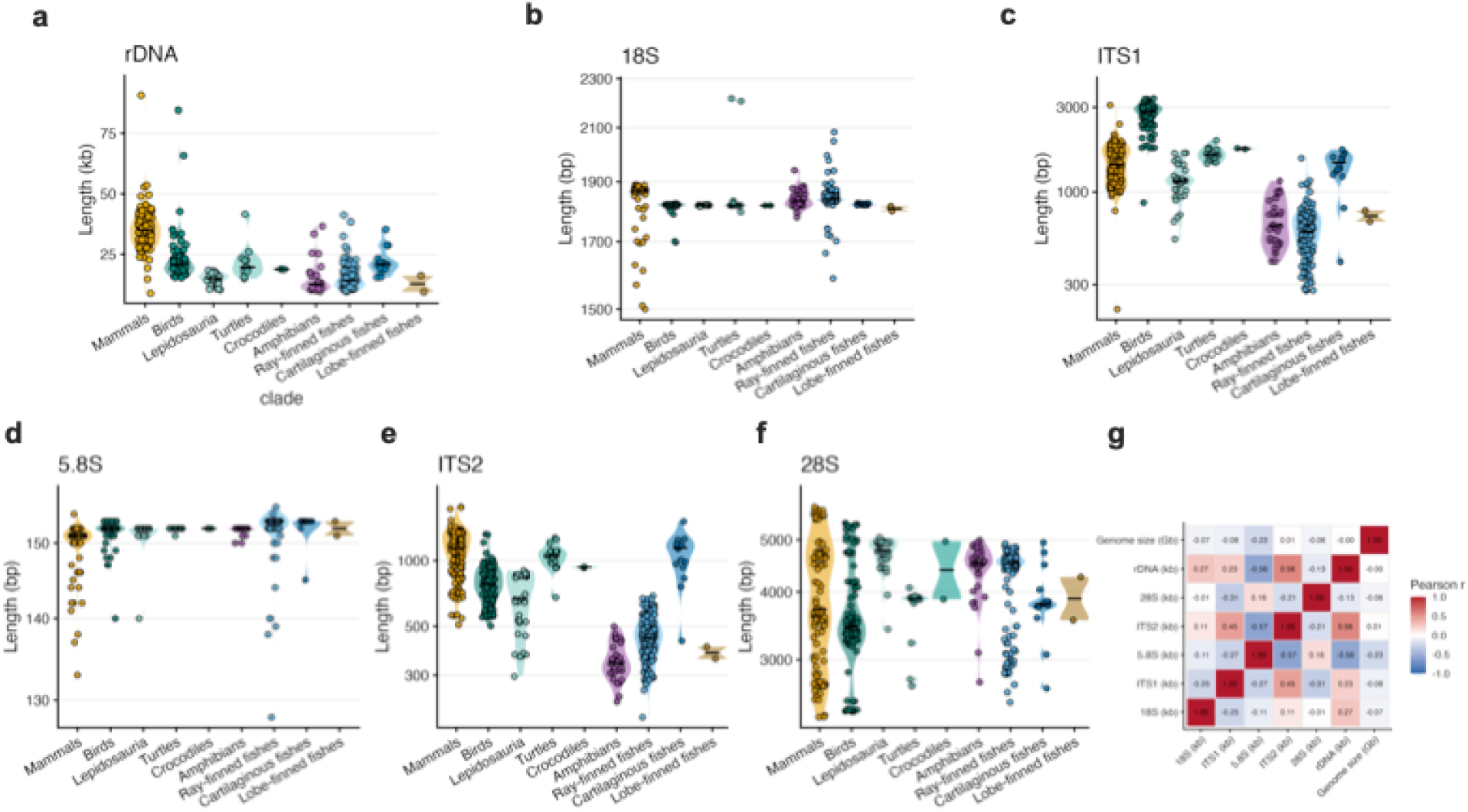
Length variation and covariation among components of the vertebrate 45S rDNA unit. (**a–f**) Distributions of sequence lengths for the full 45S rDNA unit (**a**) and its constituent regions – 18S (**b**), ITS1 (**c**), 5.8S (**d**), ITS2 (**e**) and 28S (**f)** – across major vertebrate clades. Points represent individual species and violins indicate density. Coding rRNAs (18S and 5.8S) are highly conserved in length across clades, whereas ITS1, ITS2 and 28S exhibit substantial variation, with pronounced expansion and contraction in amphibians and fishes. Total rDNA length mirrors these patterns, largely driven by spacer and 28S variability. (**g**) Pairwise Pearson correlation matrix of lengths for rDNA components and genome size. Strong positive correlations are observed between total rDNA length and ITS2 (r ≈ 0.59) and ITS1 (r ≈ 0.30), whereas coding regions show weak or negative associations with spacer lengths. Genome size is only weakly correlated with rDNA length and its components.

### rRNA sequences exhibit strikingly different patterns of length polymorphism across the vertebrate 45S rDNA repeat unit

Comparative analysis of vertebrate 45S rDNA reveals striking differences in length evolution among its constituent regions (**Fig. 3a**). Length distributions show that 5.8S rRNA is the most invariant component of the 45S unit, with nearly all vertebrate values clustering tightly around ∼150–160 bp and exhibiting minimal dispersion across clades, consistent with its extensive base-pairing interactions with 28S rRNA and its central position within the large ribosomal subunit (**Fig. 3d**). The 18S rRNA gene, which forms the scaffold of the small ribosomal subunit, also remains highly conserved but exhibits modest lineage-specific variation, typically spanning approximately 1,800–1,950 bp across vertebrates (**Fig. 3b)**. In contrast, the 28S rRNA gene shows the greatest structural flexibility among the coding rRNAs, with lengths generally ranging between roughly 3.5 kb and 5.5 kb, reflecting lineage-specific expansions and contractions of eukaryotic ribosomal expansion segments (**Fig. 3f)**. Within mammals, the 28S rRNA gene shows considerable structural stratification. Rather than forming a continuous distribution, mammalian 28S lengths segregate into four distinct clusters centered near ∼2.7 kb, ∼3.6 kb, ∼4.8 kb, and ∼5.5 kb. The intermediate cluster encompasses most placental mammals, including primates and rodents, whereas shorter 28S variants occur in several lagomorph and marsupial representatives. In contrast, the longest 28S genes are strongly enriched in chiropteran lineages, particularly vesper bats, with several species exceeding 5.6 kb. These patterns indicate that expansion segments within the large ribosomal subunit have undergone lineage-specific remodeling during mammalian evolution, producing discrete structural regimes within an otherwise highly conserved ribosomal gene.

Spacer variation is strongly structured by vertebrate phylogeny and differs between the two spacers themselves (**Supplementary Table S3**). Birds exhibit the largest ITS1 expansions, with several species exceeding 3 kb (**Fig. 3c**). Extreme examples involve ratites, including common ostrich (*Struthio camelus*), Darwin’s rhea (*Rhea pennata*), and emu (*Dromaius novaehollandiae*), all with ITS1 lengths exceeding 3.3 kb. In contrast, ITS1 values in most ray-finned fishes and amphibians remain below 1 kb. ITS2 shows a different evolutionary pattern (**Fig. 3e**). Mammals exhibit the longest ITS2 spacers, with maxima approaching ∼1.7–1.8 kb, particularly among felids such as Canada lynx (*Lynx canadensis*), jaguar (*Panthera onca*). and clouded leopard (*Neofelis nebulosa*). Amphibians and many teleost fishes show substantially shorter ITS2 regions. For example, anurans and pipids, such as *Rana temporaria*, *Bufo bufo*, and *Xenopus tropicalis*, possess ITS2 regions near the lower bound of the vertebrate distribution (∼217–230 bp), whereas several teleosts including *Danio rerio*, *Oryzias latipes*, and *Takifugu rubripes* show similarly compact spacers (∼240–260 bp). The differing clade distributions demonstrate that ITS1 and ITS2 are not simply co-expanding spacers within the same transcriptional unit but instead represent partially independent axes of rDNA evolution, influenced by lineage-specific mutational and recombinational processes.

### Spacer GC content exhibits strong phylogenetic structure

Spacer regions also show large lineage-specific differences in nucleotide composition. Median ITS1 GC content is highest in birds and lepidosaurs, whereas ray-finned fishes show the lowest values (**Fig. 4a**). Median ITS2 GC content is highest in mammals and lepidosaurs, again with ray-finned fishes occupying the lower end of the distribution (**Fig. 4b**). Extreme GC-rich spacers occur in several lineages, including the Anegada rock iguana *Cyclura pinguis* (ITS1 GC ∼0.87) and the greater mouse-tailed bat *Rhinopoma microphyllum*, which shows exceptionally high GC content in both ITS1 (∼0.86) and ITS2 (∼0.891). These findings indicate that spacer evolution involves not only length expansion but also large shifts in nucleotide composition. Phylogenetic comparison of rDNA components further illustrates the contrasting evolutionary dynamics of rRNA-encoding and spacer regions. In the circular cophylogenetic plots, the rRNA genes show relatively stable clade-level patterns consistent with the vertebrate phylogeny, whereas ITS1 and ITS2 display substantially greater heterogeneity in both length and sequence composition (**Fig. 4c-g**). This asymmetry highlights the dual evolutionary regime of the rDNA locus: deeply conserved functional genes that trace the ancient history of the ribosome, and rapidly evolving spacers that record recent lineage-specific changes in genome architecture.

**Figure 4.**
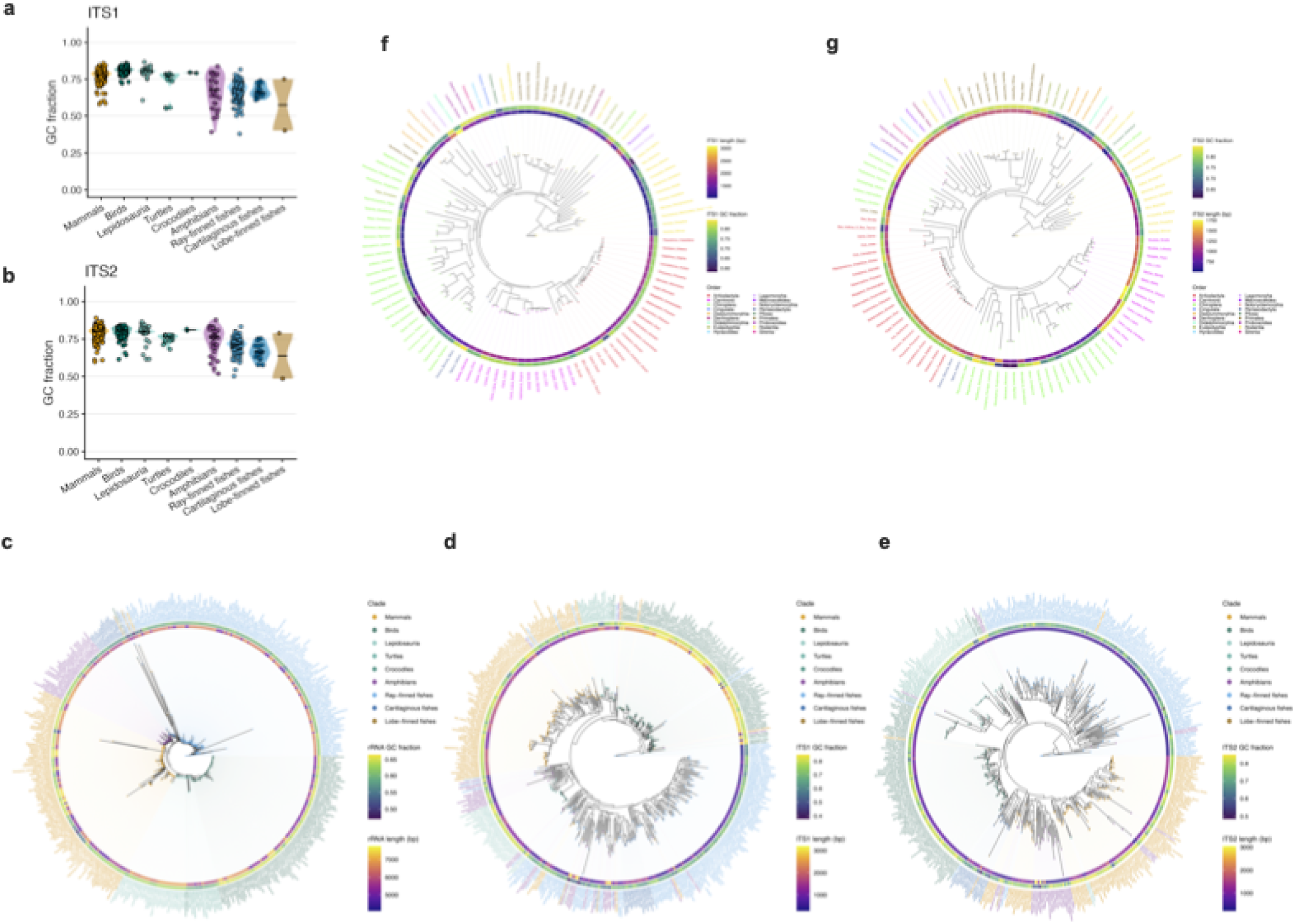
GC content variation and compositional-length covariation across 45S rDNA components and phylogeny. Violin plots showing the distribution of GC content for ITS1 (**a**) and ITS2 (**b**) across major vertebrate and selected invertebrate clades. Each point represents an individual species; violins depict density and embedded boxplots indicate median and interquartile range. (**c–g**) Circular phylogenetic trees of vertebrate taxa based on individual components of the 45S rDNA unit, with outer rings indicating sequence composition and length. Tips are colored by major clade (e.g., mammals, birds, reptiles, amphibians and fishes). For each panel, an inner ring shows GC content (continuous color scale), and an outer ring shows sequence length (bp). Panels correspond to: concatenated rRNAs (**c**), ITS1 (**d**), ITS2 (**e**), 18S (**f**) and 28S (**g**). Coding rRNAs exhibit more compact and coherent tree structure, whereas ITS1 and ITS2 show greater topological dispersion and longer branch lengths, consistent with elevated divergence and heterogeneity.

### Structure-guided alignment improves clade-level phylogenetic classification in a locus-dependent manner

Since RNA and protein structures are constrained by function, they typically evolve more slowly and retain homologous relationships longer than primary sequence alone. As a result, incorporating structural information into alignments can improve the identification of truly equivalent positions, particularly in rapidly evolving or highly divergent regions where sequence similarity becomes saturated^30,31^. Recent large-scale benchmarks show that structure-informed alignments can yield phylogenies that are more congruent with known taxonomy, especially for functional non-coding RNA, by reducing alignment noise and stabilizing homology assignment^32^. These advances motivate a direct, quantitative test of whether structural information improves phylogenetic inference across the heterogeneous components of the 45S rDNA locus. To this end, we summarized each tree by metrics quantifying how strongly taxa assigned to known clades formed contiguous blocks along the inferred topology. Each species was assigned to its expected major clade (e.g., mammals, birds, amphibians), and for each inferred tree we quantified the proportion of taxa that clustered correctly versus incorrectly within these predefined groups. This framework enables a direct, model-independent comparison of alignment strategies across all five components of the 45S rDNA locus, without requiring a fixed reference topology. Sequence divergence was estimated under the Kimura two-parameter (K2P) model, and structural divergence was measured as base-pair distance between predicted minimum-energy RNA folds.

Across loci, the impact of structure-guided alignment is strongly locus dependent (**Fig. 5a–e**). The rRNAs, particularly 18S and 28S, show high classification accuracy under both approaches, with only modest gains under structure-guided alignment. These loci already retain strong phylogenetic signal and are relatively insensitive to alignment strategy at this scale. In contrast, 5.8S – despite being the most sequence-conserved component – shows the weakest clade-level classification overall. This reflects a limitation of information content rather than alignment quality: extreme conservation stabilizes homology but provides too few informative sites to resolve deep vertebrate relationships. The clearest improvement is observed in ITS1, where structure-guided alignment substantially increases correct clade assignment and reduces fragmentation of taxa across groups (**Fig. 5b**). This indicates that ITS1 phylogenetic signal is present but partially obscured under sequence-only alignment due to homology misassignment. Incorporating secondary structure restores that signal by constraining alignment in highly variable regions. ITS2, by contrast, shows little improvement or a slight decline under the same metric (**Fig. 5d**), consistent with a higher baseline classification accuracy under sequence-only alignment and a resulting ceiling effect.

**Figure 5.**
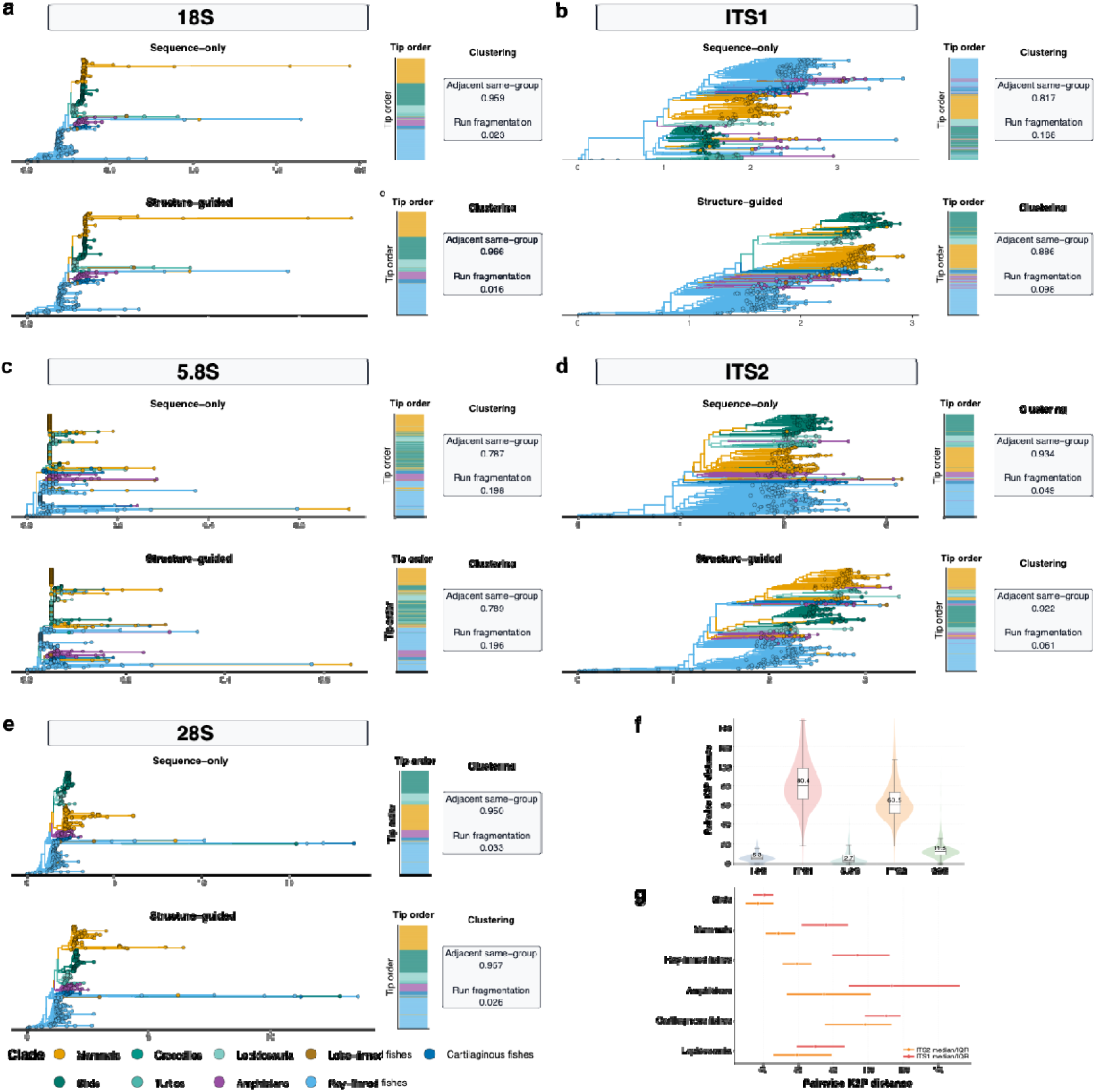
Heterogeneous phylogenetic signal across 45S rDNA loci. (**a–e**) Sequence-only and structure-guided phylogenetic trees for 18S (**a**), ITS1 (**b**), 5.8S (**c**), ITS2 (**d**), and 28S (**e**), respectively. For each locus, the upper row shows the sequence-only tree and the lower row shows the structure-guided tree. Branches and terminal tips are colored by clade, and the adjacent vertical strip summarizes the same clade assignment in terminal tip order. Two tip-order clustering metrics were reported to each tree: the adjacent same-group fraction, where larger values indicate that neighboring terminal taxa more frequently belong to the same clade, and the run fragmentation index, where smaller values indicate fewer interruptions of same-clade blocks along the terminal order. (**f**) Distribution of pairwise sequence divergence (Kimura two-parameter, K2P) across loci. Coding rRNAs (18S and 5.8S) show low divergence with narrow distributions, whereas ITS1 exhibits the highest divergence and widest spread (median ∼80), followed by ITS2 (∼60). The 28S gene shows intermediate divergence (∼12), higher than 18S but far below spacer regions. (**g**) Within-clade divergence for ITS1 and ITS2 (median ± IQR). Across all major clades, ITS1 consistently shows higher divergence than ITS2, with the largest values observed in amphibians and cartilaginous fishes, and comparatively reduced divergence in birds.

These classification results map directly onto the divergence landscape of the 45S locus (**Fig. 5f–g**). Coding rRNAs exhibit low sequence divergence and consistent clustering, whereas ITS1 and ITS2 show elevated divergence and pronounced heterogeneity across clades. ITS1 emerges as the most labile component, with high sensitivity to alignment strategy, while ITS2 retains stronger baseline clustering but exhibits reduced responsiveness to structural information. At shallow phylogenetic scales, both spacers recover taxonomic groupings, but their signal progressively degrades with evolutionary depth.

This decoupling is further exemplified by the ITS1–ITS2 cophylogeny within mammals (**Fig. 6**), where systematic crossing patterns reveal that the two spacers do not represent noisy versions of a single signal but instead follow distinct evolutionary trajectories within the same rDNA array. For example, Chiroptera show pronounced internal reshuffling between ITS1 and ITS2 despite clear taxonomic cohesion, indicating differential spacer evolution within the same lineage. Rodents similarly cluster more tightly in ITS1 but fragment in ITS2, highlighting asymmetry in divergence patterns even among relatively close taxa. Artiodactyls (including cetaceans) broadly retain their grouping but display partial breakdown of expected sister relationships, and carnivores exhibit subtler “micro-rearrangements” that preserve higher-order structure while altering fine-scale topology. Occasional cross-order link crossings further point to homoplasy or limits of alignment homology in highly divergent spacers.

**Figure 6.**
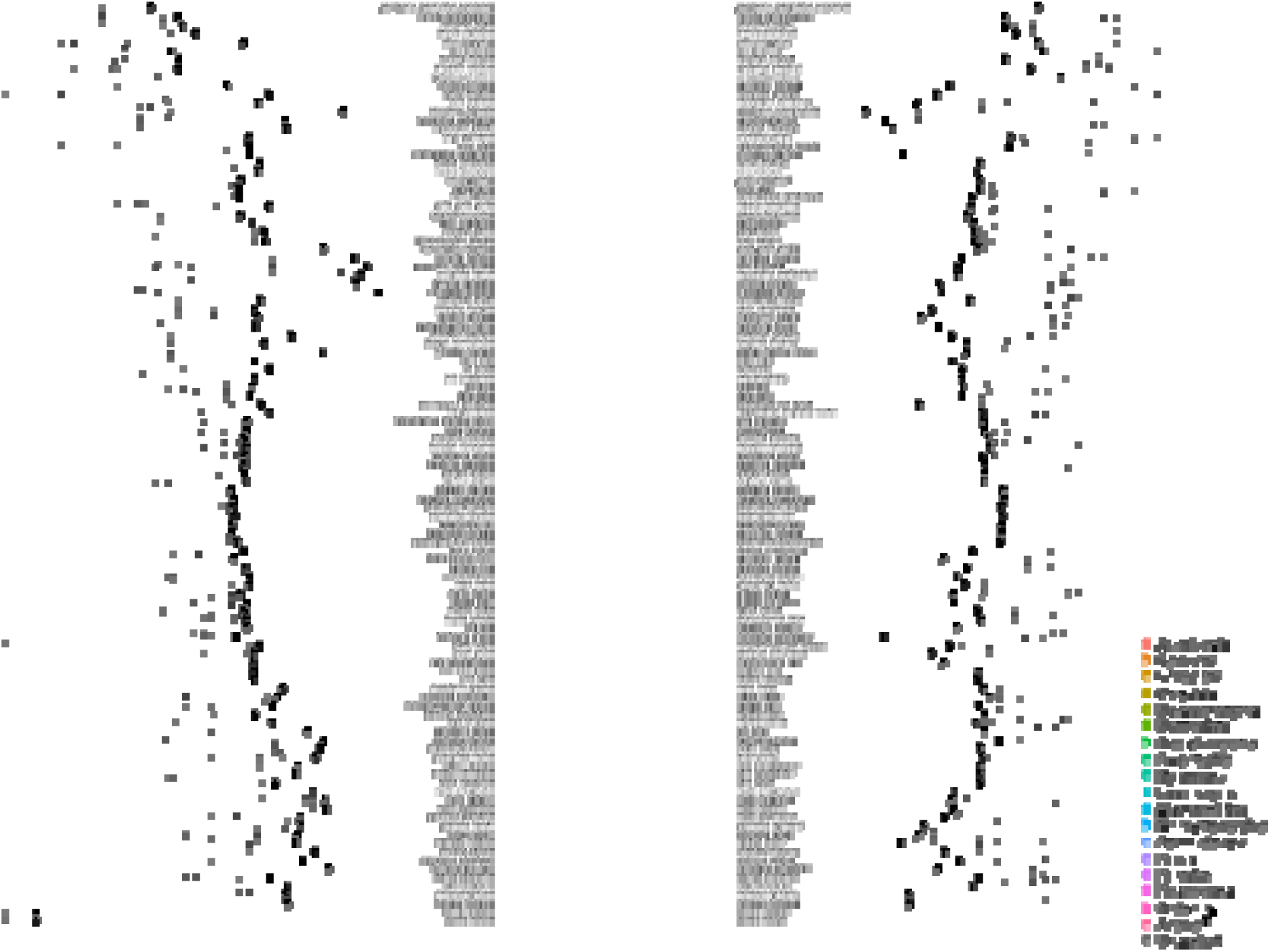
Tanglegram comparing mammalian phylogenies derived from ITS1 and ITS2 regions. Lines connect identical taxa across trees, illustrating substantial topological discordance between spacer-based and coding-gene phylogenies, with extensive crossing indicating reduced phylogenetic congruence of ITS sequences.

These patterns indicate that ITS1 and ITS2 are not simply noisy versions of the same signal, but differentially evolving markers shaped by locus-specific rates, structural constraints, and the stochastic dynamics of concerted evolution within multi-copy rDNA arrays. This supports a layered model of 45S evolution in which 18S and 5.8S behave as highly conserved phylogenetic anchors, 28S occupies an intermediate position, and ITS1/ITS2 represent rapidly diverging, only partially phylogeny-faithful regions whose signal increasingly reflects lineage-specific turnover and homology breakdown at deeper timescales.

### Sequence and secondary structure coevolution

To further quantify the relationship between sequence and structural evolution across the 45S rDNA unit, we analyzed pairwise divergence among the three rRNA coding genes (18S, 5.8S, and 28S) and the internal transcribed spacers (ITS1 and ITS2) across vertebrates using a unified computational framework. The small and intermediate rRNA components (18S and 5.8S) exhibited the strongest coupling between sequence and structure (**Fig. 7**). Pairwise sequence divergence remained low and tightly distributed, typically below ∼10–15 substitutions per 100 sites and was strongly correlated with structural divergence (Pearson’s r = 0.61 and 0.57, respectively; Mantel r ≈ 0.6, p = 0.001). Despite spanning deep evolutionary distances, these loci show striking conservation of both sequence and folding architecture. For example, 5.8S rRNA sequences from phylogenetically distant taxa such as *Abramis brama* (ray-finned fish), *Alligator mississippiensis* (crocodilian), and *Balaenoptera musculus* (mammal) remain nearly identical in length and predicted secondary structure, with only minor compensatory substitutions preserving canonical stem–loop interactions. Similarly, 18S rRNA exhibits highly conserved global folding across vertebrates, with variation largely confined to small peripheral regions that do not disrupt the core ribosomal scaffold.

**Figure 7.**
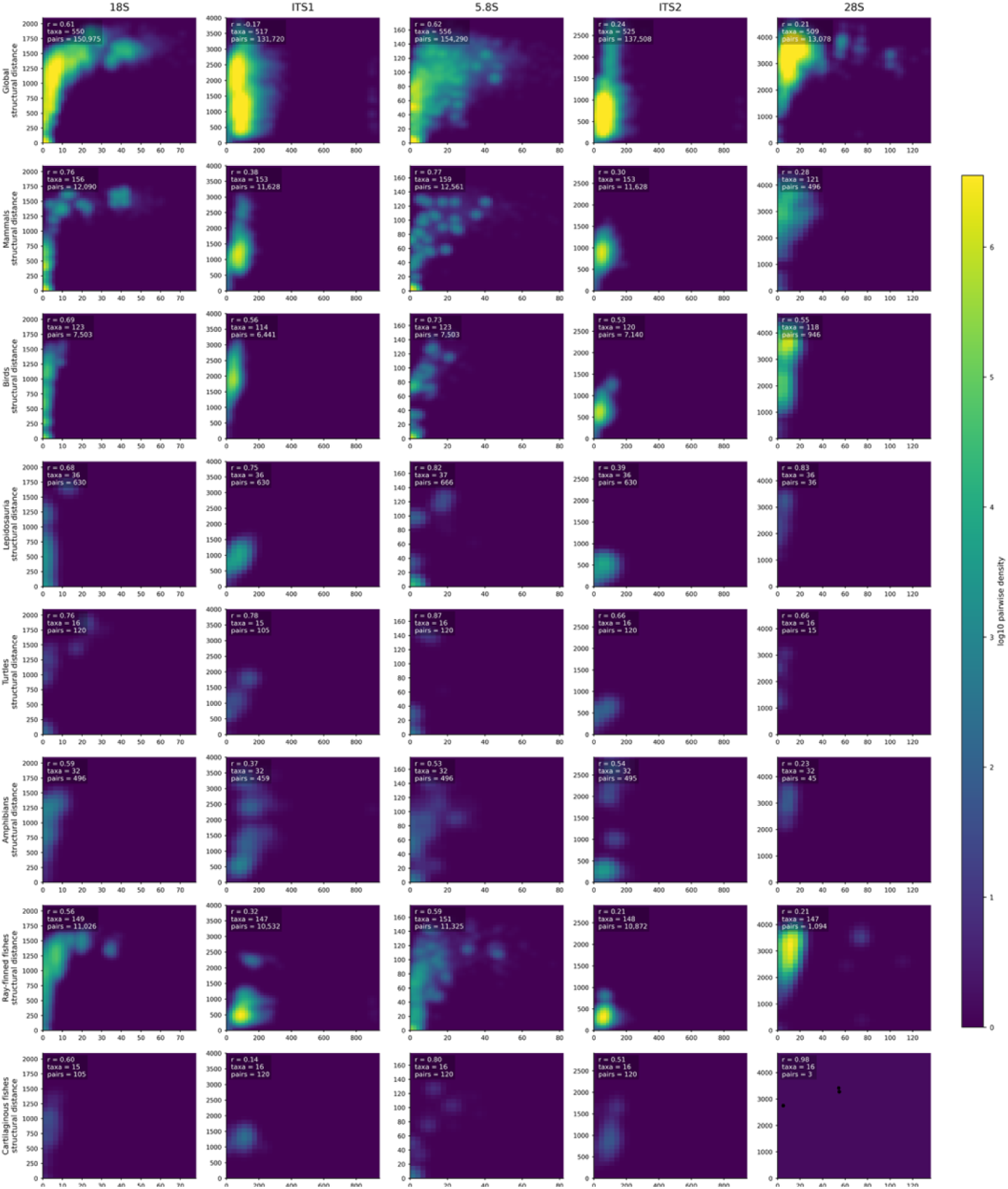
Clade-specific sequence–structure landscapes across the vertebrate 45S rDNA unit. Pairwise relationships between sequence divergence (K2P) and RNA secondary-structure divergence are shown for 18S, ITS1, 5.8S, ITS2, and 28S (columns). The top row (“Global”) includes all vertebrates; subsequent rows show within-clade comparisons. Panels are rendered as two-dimensional density heat maps (log-scaled), with color indicating the local density of pairwise comparisons; sparse datasets are shown as points. Insets report Pearson’s correlation (r), number of taxa, and number of pairwise comparisons. The coding rRNAs (18S and especially 5.8S) exhibit strong and consistent coupling between sequence and structure, forming compact, diagonal density distributions across clades. In contrast, ITS1 and ITS2 display broader and more heterogeneous landscapes, indicating weaker and more variable coupling, consistent with modular spacer evolution. The 28S rRNA shows comparatively weak and asymmetric relationships, reflecting its composite architecture with lineage-specific expansion segments. Together, these patterns highlight a conserved sequence–structure regime in core rRNAs versus a more flexible, clade-dependent evolutionary landscape in the spacers.

The large subunit rRNA (28S) showed a distinct pattern. Although structurally complex and functionally constrained, 28S showed near-zero correlation between sequence and structural divergence (Pearson’s r ≈ 0.002; Mantel r ≈ −0.002). This apparent decoupling likely reflects saturation effects combined with lineage-specific expansion segments. For instance, comparisons between amphibians such as *Ascaphus truei* and birds such as *Aythya fuligula* reveal substantial sequence divergence concentrated in expansion segments, while the conserved core helices remain structurally stable.

### ITS2: modular evolution under partial structural constraint

ITS2 sequences show extensive variation in both sequence length and nucleotide composition across vertebrate clades, ranging from ∼300 bp in many teleost fishes to >1500 bp in several passerine birds and mammals. Sequence divergence within major vertebrate clades is moderate but clearly structured phylogenetically. Mammalian ITS2 sequences cluster into a relatively cohesive group with moderately long spacers (∼900–1000 bp) and high GC content. Despite this dramatic size heterogeneity, predicted secondary structures retain a broadly conserved architecture characterized by a multi-helix scaffold radiating from a central loop, consistent with the canonical ITS2 folding model described across eukaryotes. Several rodent lineages, including *Acomys* species, exhibit particularly expanded ITS2 regions approaching ∼1 kb, accompanied by extensive internal stem-loop structures with predicted folding energies exceeding −500 kcal/mol. In contrast, many ray-finned fishes possess substantially shorter ITS2 spacers (∼300–500 bp) and correspondingly simpler secondary structures with fewer long helices. For example, cypriniform fishes such as *Abramis brama* display compact ITS2 structures of ∼300 bp with predicted minimum free energies near −160 kcal/mol, whereas sturgeons (*Acipenser*) show intermediate lengths (∼530 bp) but maintain relatively stable multi-helix folds. Birds represent one of the most structurally elaborate ITS2 groups in the dataset. Several passerines show long ITS2 spacers exceeding 900 bp, frequently dominated by GC-rich sequence tracts and repetitive elements that generate large, highly stable secondary structures with folding energies below −600 kcal/mol. In extreme cases, such as *Agelaius phoeniceus*, ITS2 length exceeds 1.5 kb, producing exceptionally complex predicted structures with numerous nested helices. Crocodilians and other reptiles generally show intermediate ITS2 sizes (∼800–950 bp) with stable multi-helix structures broadly similar to those seen in mammals.

Importantly, sequence divergence and secondary-structure complexity are not strictly proportional. Some taxa with modest ITS2 lengths exhibit highly stable folds due to elevated GC content, whereas other taxa with longer spacers accumulate expansion segments that contribute limited additional base-paired structure. Across 137,550 pairwise comparisons, sequence divergence and structural divergence were positively but only weakly correlated (Pearson’s r = 0.215; Spearman’s ρ = 0.110), indicating that substantial sequence divergence can accumulate without proportional structural divergence. Because raw base-pair distance is influenced by spacer length, we also normalized structural distance by the mean ITS2 length of each species pair; the relationship remained weakly positive (r ≈ 0.194), confirming that the signal is not driven solely by absolute spacer size. Given the non-independence of pairwise comparisons, we additionally performed Mantel tests (r = 0.283, p = 0.001). The strength of sequence–structure coupling varied among clades, being moderate in amphibians, birds, mammals, and lepidosaurs, but weak in ray-finned fishes and nearly absent in cartilaginous fishes.

ITS2 sequences retain a conserved secondary structure characterized by a core scaffold of helices II and III (**Fig. 8**). These domains contain the most conserved primary sequence motifs and maintain stable base-pairing interactions across vertebrates. In contrast, peripheral helices (I and IV) and the extended 3′ domain exhibit substantial lineage-specific expansion and structural diversification. This modular organization explains the weak but significant correlation between sequence and structural divergence, indicating that ITS2 evolution is constrained at the level of global architecture while allowing extensive local sequence and structural plasticity. Together, these results indicate that vertebrate ITS2 evolves under a regime of rapid primary-sequence turnover coupled to more gradual structural divergence, consistent with preservation of a constrained RNA scaffold despite extensive nucleotide change. Such patterns are consistent with the concept of compensatory base changes maintaining ITS2 structural integrity despite rapid sequence turnover. Across vertebrates, therefore, ITS2 evolution appears to follow a dual regime: rapid primary sequence divergence combined with partial structural conservation of the underlying RNA fold. This combination likely reflects the functional constraints imposed by ITS2’s role in pre-rRNA processing and ribosome assembly, where secondary structure rather than primary sequence provides the key functional scaffold.

**Figure 8.**
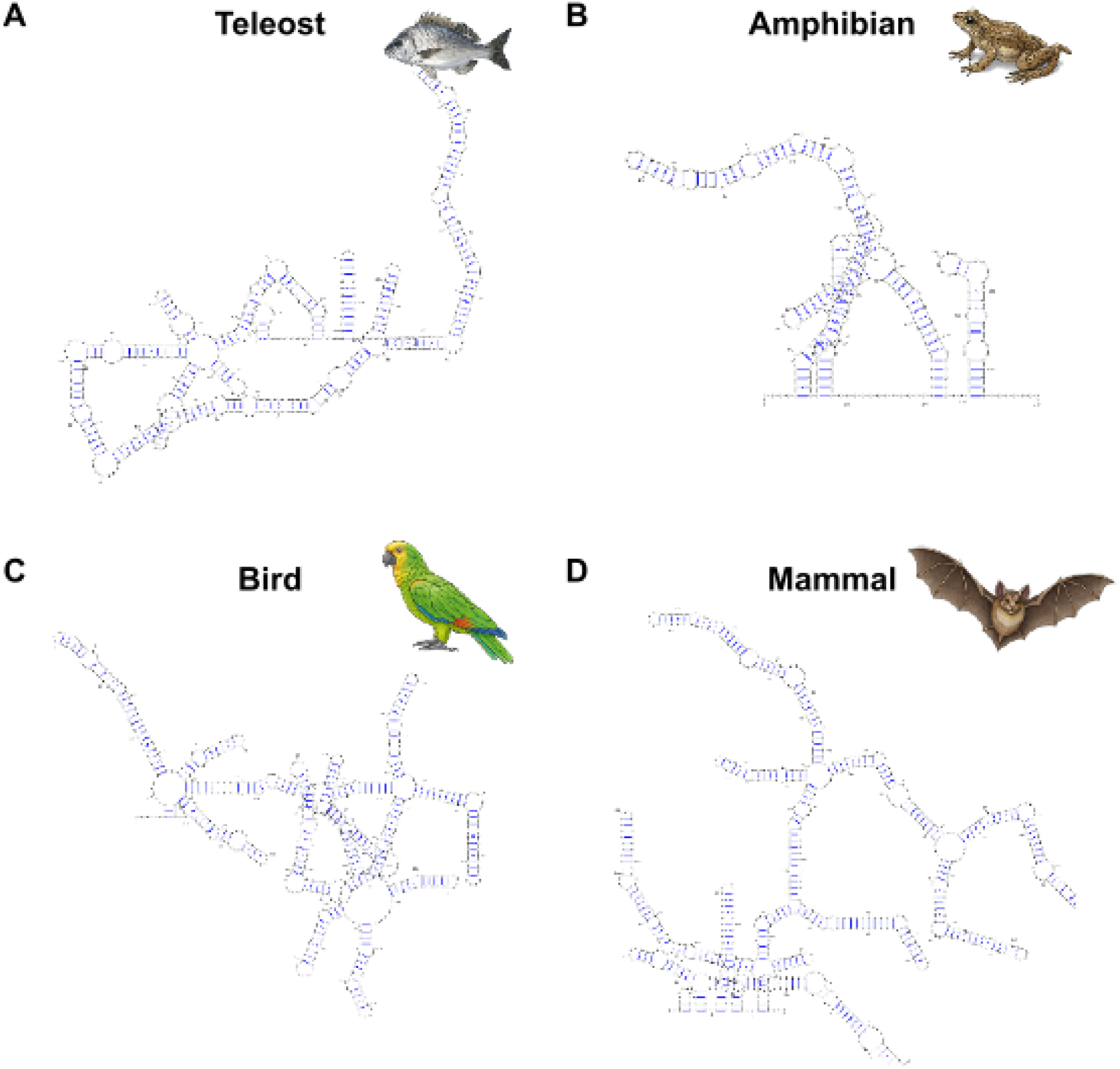
Comparative secondary structure of ITS2 across major vertebrate clades. Predicted minimum free-energy secondary structures of the internal transcribed spacer 2 (ITS2) are shown for representative species from four vertebrate lineages: (A) teleost fish (*Acanthopagrus schlegelii*), (B) amphibian (*Ascaphus truei*), (C) bird (*Amazona ochrocephala*), and (D) mammal (*Artibeus lituratus*). Structures were inferred using thermodynamic folding algorithms and visualized in standardized layouts to facilitate comparison. Base-paired regions are indicated by canonical Watson–Crick interactions, with helices radiating from a central core domain. Despite substantial sequence divergence, the overall architectural organization of ITS2, including a conserved central ring and multiple radiating helices, is maintained across taxa, with lineage-specific variation in helix length and branching complexity.

### ITS1: breakdown of global structural constraint

ITS1 represents the least constrained component of the 45S rDNA unit. Sequence divergence frequently reaches extreme values exceeding 100–200 substitutions per 100 sites accompanied by dramatic length expansion. For example, anseriform birds such as *Anas platyrhynchos* and passerines such as *Acridotheres tristis* routinely exhibit ITS1 lengths approaching ∼3 kb, whereas amphibians such as *Ambystoma mexicanum* show similarly extreme expansion. In contrast, basal ray-finned fishes such as *Amia calva* retain compact ITS1 regions of only a few hundred base pairs, illustrating a >10-fold length disparity across vertebrates. Structurally, ITS1 lacks a conserved global architecture. Predicted folds in compact fish ITS1 sequences show relatively simple stem–loop arrangements, whereas expanded mammalian (e.g., *Acomys minous*) and avian ITS1 sequences form deeply nested and highly complex structures with extensive long-range pairing. These structures are often non-homologous across taxa and lack recognizable shared domains. Consistent with this heterogeneity, sequence and structural divergence are negatively correlated (Pearson’s r ≈ −0.17; Spearman’s ρ ≈ −0.30), and many pairwise structural comparisons are non-comparable, reflecting breakdown of shared folding architectures. Together, these observations indicate that ITS1 evolves under a regime of extreme evolutionary flexibility, with rapid sequence turnover and weak constraints on global structure.

### Thermodynamic scaling of RNA stability across the 45S rDNA unit

These structural regimes are further reflected in the thermodynamic stability of predicted RNA folds. The rRNA coding genes exhibit relatively constrained folding energies consistent with their conserved size and architecture. In contrast, ITS regions span a much broader energetic range, scaling with both sequence length and GC content. Compact ITS2 structures in fishes such as *Alosa sapidissima* show folding energies near −150 kcal/mol, whereas large avian or reptilian ITS2 sequences (e.g., *Anolis sagrei*) can drop below −600 kcal/mol. ITS1 extends this trend further, with long, GC-rich spacers producing extremely stable but structurally heterogeneous folds with predicted energies approaching −500 kcal/mol or lower despite lack of conserved architecture, exemplified by mammals, such as *Apodemus sylvaticus*. Importantly, increased thermodynamic stability does not necessarily correspond to conserved structural organization. Instead, highly negative folding energies in ITS1 and ITS2 often reflect redundant or non-specific base pairing within expanded sequence tracts, reinforcing the conclusion that sequence divergence can greatly exceed meaningful structural divergence in spacer regions.

### A continuum of evolutionary regimes within the 45S rDNA unit

Taken together, these results reveal a hierarchical organization of evolutionary constraint within the 45S rDNA repeat. The rRNA coding genes (18S and 5.8S, and to a lesser extent 28S) occupy a regime of strong sequence–structure coupling driven by functional constraints on ribosome architecture. ITS2 represents an intermediate regime in which sequence divergence exceeds structural divergence but remains anchored by a conserved RNA scaffold. ITS1, in contrast, approaches a regime of structural instability and non-comparability, reflecting rapid evolutionary turnover unconstrained by a fixed global architecture. This gradient demonstrates that rDNA evolution is not governed by a single set of constraints but instead reflects a continuum of evolutionary dynamics operating within a single transcriptional unit – from deeply conserved structural RNAs to rapidly evolving spacer elements with progressively relaxed structural constraints.

## Discussion

The Vertebrate Genomes Project (VGP^25^) provides, for the first time, a phylogenetically broad and assembly-resolved view of the 45S rDNA locus, allowing long-standing cytogenetic observations to be re-evaluated in a genomic framework. What emerges from this analysis is not a static or purely housekeeping system, but a dynamic genomic compartment in which deep sequence conservation, divergence, and structural plasticity coexist. Across vertebrates, the 45S rDNA locus retains a remarkably stable functional core, yet varies extensively in copy number, chromosomal position, and spacer architecture. This combination of stability and flexibility has been noted in isolated systems, but the present dataset demonstrates that it is a general property of vertebrate genomes rather than an exception.

The chromosomal organization of rDNA reinforces this pattern. While a tendency toward distal, often subtelomeric localization is evident, particularly in mammals and many fishes, this pattern is neither universal nor rigid. Instead, vertebrate rDNA occupies a spectrum, from relatively constrained architectures in amniotes to markedly dispersed configurations in amphibians and certain fish lineages ^10^. Such variability is consistent with earlier cytogenetic surveys showing that rDNA loci can occur at nearly any chromosomal position, despite statistical biases toward terminal or pericentromeric regions^27^. The VGP data extend this by placing these distributions into a phylogenetic context, suggesting that chromosomal positioning is shaped not only by mechanistic constraints, such as recombination or chromatin organization, but also by lineage-specific evolutionary trajectories. In this sense, rDNA loci behave less like fixed genomic landmarks and more like mobile elements embedded within broader chromosomal dynamics.

Copy number variation adds another layer to this picture. The range of rDNA repeat numbers observed across eukaryotes is extreme, spanning orders of magnitude, yet only a subset of these copies is transcriptionally active at any given time^14,33,34^ . This decoupling of copy number from functional output complicates simple interpretations of rDNA abundance. Rather than reflecting transcriptional demand alone, copy number appears to represent a form of genomic capacity, maintained through recombination-driven turnover and balanced by mechanisms that restore repeat number when it declines^35^. The variability observed across vertebrates is therefore not incidental but intrinsic to the system, consistent with the idea that rDNA arrays function as a flexible genomic reservoir with roles extending beyond ribosome production, including genome stability and global regulation of gene expression^29^. Previous cytogenetic surveys have highlighted the remarkable variability in rDNA copy number and chromosomal positioning across animals ^27^. Our results extend this framework by resolving the internal evolutionary dynamics of the 45S locus, revealing that the apparent coherence of rDNA at the chromosomal level masks a pronounced heterogeneity in sequence–structure coupling and evolutionary constraint across its constituent elements.

Underlying both the dynamic copy-number variation and the apparent coherence of rDNA repeats is the process of concerted evolution, long understood to arise from a combination of unequal crossing over and gene conversion^5^, two mechanistically related outcomes of homologous recombination. These processes act to homogenize repeats within arrays over evolutionary time, maintaining high sequence similarity despite rapid turnover. However, recombination is not necessarily composition-neutral. Recombination can be associated with GC-biased fixation processes, including GC-biased gene conversion (gBGC), in which repair of mismatches during recombination preferentially favors G/C over A/T alleles^36,37^. Because rDNA arrays are among the most recombinogenic regions of the genome, they provide a natural context in which such biases can accumulate over time. Thus, the same molecular mechanisms that drive concerted evolution and maintain repeat homogeneity may also contribute to shaping base composition, linking recombination dynamics to the elevated and lineage-variable GC content observed across the 45S locus. Our results are broadly consistent with that interpretation at the level of overall composition: the elevated GC content of vertebrate 45S rDNA cannot be explained by unequal crossing over alone, which is composition-neutral, and instead points to GC-biased fixation processes such as gBGC^36^. However, the expanded phylogenomic scope reveals a critical limitation of the earlier proxy-based view. Rather than exhibiting a coherent compositional signal, the 45S array is partitioned into components with distinct GC regimes, such that GC enrichment in one region cannot be assumed to represent the entire repeat. This pronounced heterogeneity indicates that any GC-biased process operates in a context-dependent manner, interacting with local functional and structural constraints, strongly constraining the rRNAs while permitting more variable equilibria in the spacers, rather than imposing a uniform compositional landscape across the locus. In this framework, concerted evolution homogenizes repeats within regions but does not override the establishment of region-specific GC equilibria.

Similarly, at the level of sequence organization, the 45S rDNA locus emerges not as a single evolutionary unit, but as a composite system in which fundamentally different regimes of constraint coexist within a contiguous transcriptional framework. Across vertebrates, the rRNAs, 18S and 5.8S in particular, exhibit strong phylogenetic signal, low sequence divergence, and tight coupling between sequence and secondary structure, consistent with the extensive functional constraints imposed by ribosome assembly and translational fidelity. These observations align with decades of comparative work showing that small subunit rRNA evolves under intense purifying selection, maintaining both sequence identity and higher-order folding across deep evolutionary timescales^3^. 5.8S rRNA represents a eukaryote-specific addition to the ribosome and the reason for this evolutionary novelty is unknown. Its pronounced conservation likely reflects an essential role in stabilizing interactions within the large subunit, including contacts with 28S rRNA, and possibly in facilitating ribosome dynamics during translation^38^. Consistent with this functional importance, mutations in 5.8S rRNA impair cell growth and protein synthesis, and are associated with aberrant ribosomal profiles, including enlarged polyribosomes and altered ribosome-associated tRNA occupancy^38^.

The large subunit 28S rRNA occupies an intermediate position within this framework. Although it remains a coding component of the ribosome, its sequence–structure coupling is weaker and its divergence more pronounced than in 18S and 5.8S. This pattern is consistent with the modular organization of 28S, which contains both highly conserved core domains and rapidly evolving expansion segments^39^. These expansion regions are known to vary extensively in length and structure across eukaryotes and are thought to contribute to lineage-specific ribosomal features^40^. Their relative freedom from strict functional constraint likely underlies the broader dispersion observed here, indicating that even within coding rRNAs, evolutionary pressures are unevenly distributed.

In contrast, the internal transcribed spacers display a markedly different evolutionary profile. Both ITS1 and ITS2 show elevated sequence divergence and pronounced heterogeneity across clades. Within the spacers themselves, a further asymmetry becomes apparent. ITS2 retains a recognizable structural framework across taxa, despite substantial sequence divergence, whereas ITS1 appears more labile and less constrained. This pattern is broadly consistent with their functional role, which is largely associated with pre-rRNA processing and does not extend to the mature ribosome. As a consequence, selection acts indirectly, primarily through constraints on RNA folding and processing efficiency rather than precise nucleotide identity^41,42^. The resulting relaxation of sequence constraint allows spacers to accumulate substitutions, insertions and deletions at rates far exceeding those observed in coding rRNAs, while still retaining the minimal structural features required for correct maturation. In ITS2 in particular, conserved architectural elements, including the canonical multi-helix folding pattern, can be maintained despite extensive sequence turnover, consistent with previous studies demonstrating that selection acts on structural motifs rather than exact sequence composition^21,43^. Notably, spacer transcript fragments of ITS2 may also be co-opted into RIG-I signaling pathway, contributing to the sensing of pathogen-derived nucleic acids^44^. The results reinforce the view that rDNA cannot be understood solely in terms of ribosome biogenesis. The nucleolus, organized around rDNA arrays, is being increasingly recognized as a multifunctional hub involved in processes ranging from cell cycle regulation to stress responses^10^, and the genomic behavior of rDNA reflects this expanded role. Variation in copy number, chromosomal position, and spacer composition is therefore likely to have consequences that extend beyond translation, influencing genome organization and regulatory landscapes at multiple levels.

A central feature of our results is the gradient in sequence–structure coupling across the 45S rDNA locus. In 18S and 5.8S, sequence and structural divergence are strongly correlated, reflecting the well-established principle of compensatory evolution in paired RNA regions, whereby mutations disrupting base pairing are offset by coordinated substitutions that restore structural integrity. The persistence of this relationship across hundreds of vertebrate taxa indicates that the underlying constraint architecture of ribosomal RNAs remains stable even over deep phylogenetic timescales. By contrast, the weaker coupling observed in ITS regions suggests that structural conservation operates within a broader sequence space. This decoupling is reflected in the asymmetry between sequence and structural divergence in spacer regions, where sequence distances consistently exceed structural distances. Such behavior is expected under models of RNA evolution in which multiple distinct sequences can adopt similar minimum-energy folds, effectively buffering structural features against rapid sequence change. This many-to-one mapping between sequence and structure has been described as a general property of RNA genotype–phenotype landscapes^45^, and our results extend its relevance to large-scale comparative analyses of vertebrate rDNA. The implication is that structural conservation in ITS regions represents a lower-dimensional constraint that permits substantial evolutionary flexibility at the sequence level, while still preserving essential aspects of RNA folding required for processing.

At the locus level, the contrasting behaviors of coding and spacer regions are also consistent with differential operation of concerted evolution. Mechanisms such as unequal crossing-over and gene conversion act to homogenize rDNA repeats within genomes, thereby maintaining low intra-species variation in conserved regions^35^. The greater heterogeneity observed in ITS sequences suggests either reduced efficiency of these processes or a balance between homogenization and rapid sequence turnover. This interplay likely contributes to the maintenance of functional coherence within rDNA arrays while permitting diversification in less constrained regions.

Our findings have also direct implications for the use of rDNA in phylogenetics. The strong conservation and high sequence–structure coupling of 18S and 5.8S reinforce their suitability for resolving deep evolutionary relationships, whereas the rapid evolution of ITS regions supports their continued use as markers of recent divergence. However, the observed decoupling between sequence and structure in spacers also suggests that phylogenetic inference based solely on sequence data may overlook informative structural constraints, particularly in highly divergent datasets. Previous work has shown that incorporating ITS2 secondary structure can improve alignment accuracy and phylogenetic resolution^46^, and our results provide a quantitative framework for understanding why such approaches are effective. The present comparison, spanning hundreds of vertebrate genomes and multiple rDNA components analyzed in parallel, provides an unprecedented framework for evaluating methodological performance. Structure-informed phylogenies turn out to show modest but consistent improvements in clade coherence relative to sequence-only approaches, with the magnitude of the effect varying across the 45S locus. Even though incorporating secondary structure does not radically alter phylogenetic inference, it can improve alignment stability and slightly reduce misassignment, especially in more divergent comparisons. Thus, rather than representing a categorical shift in performance, structure-informed approaches provide incremental gains that become more relevant as sequence signal degrades.

By spanning the full range from LUCA-scale molecular conservation to rapid lineage-specific divergence, ribosomal DNA encapsulates a dual tempo of genome evolution within a single locus. Beyond their importance for phylogenetics, these results underscore the necessity of resolving rDNA repeats to achieve truly complete and biologically meaningful reference genomes and establish vertebrate rDNA as a powerful system for studying concerted evolution, genome organization, and the evolutionary dynamics of essential cellular machinery. As long-read genomics continues to close the final gaps in vertebrate assemblies, rDNA emerges not as an assembly nuisance, but as a uniquely informative substrate for evolutionary genomics. In this context, the contribution of the VGP is not merely technical but conceptual. By resolving rDNA loci across a wide phylogenetic range, it allows patterns that were previously inferred from fragmented data to be observed directly and comparatively.

## Methods

### Vertebrate genome assemblies

We analyzed chromosome-level or near chromosome-level assemblies from VGP Phase I, available via assembly hub at UCSC (hgdownload.soe.ucsc.edu/hubs/VGP/). The VGP data freeze ^25^ covers all major vertebrate clades including mammals, birds, reptiles, amphibians, ray-finned and cartilaginous fishes, cyclostomes, and other deuterostomes.

### 45S rDNA identification and annotation

Assembled genomes were scanned for nuclear rDNA with barrnap^47^ (kingdom = eukaryote) to locate 5.8S rRNA loci. For each 5.8S rRNA hit, a flanking 15 kb genomic window (30 kb total) was extracted, and sequences were retrieved with BEDTools getfasta^48^. Internal transcribed spacers (ITSs) were identified with ITSx^49^ using ‘taxon = Metazoa’, the ‘-only_full’ option, and saving ITS1 and ITS2 regions. Only full ITS1 predictions delimited by SSU-5.8S and full ITS2 predictions delimited by 5.8S-LSU were retained. Quality control filters were applied per ITS sequence. We computed the fraction of ambiguous bases (N) and the maximum homopolymer run length; sequences with N-fraction > 0.05 or any homopolymer > 15 nt were removed. For each species and locus (ITS1 or ITS2), we summarized the retained copies into per-species statistics. 45S rRNA transcription units (18S, 5.8S, 28S) were annotated directly on assemblies using barrnap^47^ with the eukaryotic model. Partial predictions were excluded. For every species and each rRNA type, all full-length features were collected and used to extract the corresponding nucleotide sequences from the genome assembly. Within each species and rRNA unit, a single representative copy was chosen rather than constructing a multiple-sequence consensus. For a given unit, the lengths of all valid copies were summarized, and the median length was computed. The representative rRNA gene copy was then defined as the sequence whose length was closest to this species-specific median, which was used as input for downstream multiple sequence alignment and phylogenetic analyses.

### Consensus ITS sequences

For each species and locus (ITS1 or ITS2), we collected all high-quality ITS predictions passing the filters described above. Within each species-locus combination, sequences were collapsed to unique haplotypes and clustered at high pairwise identity (98%) to group near-identical copies into within-species ITS clusters. For the consensus sequence, only sequences belonging to the largest cluster for that species were aligned and a majority-rule consensus sequence was called, using standard IUPAC ambiguity symbols where no single base reached a strict majority at a given position. Per-species consensus sequences were served as the input for downstream multiple-sequence alignment and phylogenetic analyses.

### rDNA cluster size estimation

To estimate the total size of complete rDNA transcriptional clusters directly from genome assemblies, we parsed the same barrnap GFF files for each species and reconstructed 45S arrays on each contig and strand. For each contig/strand combination we: 1) Collected all rRNA hits (18S, 5.8S, and effective 28S) and sorted them by genomic coordinate. 2) Identified the ordered list of 18S start coordinates. 3) For each consecutive 18S pair, we defined an interval from the start of one 18S to the start of next 18S. 4) Within this interval we counted occurrences of 5.8S and 28S. When at least one 5.8S and one 28S occurred between the two 18S starts, we defined a complete rDNA cluster. The rDNA repeat unit length was defined as the distance between the two 18S starts (inclusive). To guard against spurious long intervals due to assembly gaps or misannotations, we only retained units with length ≤ 100 kb, consistent with and extending beyond the ∼50 kb upper range reported in mammals^50^. To assess the performance of the pipeline, we reanalyzed the complete human T2T-CHM13 assembly^6^ using the same workflow. The pipeline recovered 219 complete 45S rDNA copies (Supplementary Table S3) spanning 9.9 Mb of assembled sequence, identical to the values reported for T2T-CHM13^6^. This agreement supports the ability of our approach to recover repeat-unit content and overall array size in a benchmark genome with extensively characterized rDNA arrays. Nevertheless, inferred copy numbers across species remain dependent on assembly completeness and should be regarded as approximate estimates of assembled rDNA content.

### Phylogenetic reconstruction

For phylogenetic analyses, we generated multiple sequence alignments separately for 18S, 5.8S, ITS1, ITS2, 28S consensus sequences, and 45S rRNA (concatenated 18S, 5.8S, and 28S) using MAFFT^51^. For sequence-only alignments, loci expected to be globally alignable, 18S and 5.8S, were aligned with the G-INS-i strategy using ‘--globalpair --maxiterate 1000’, whereas more variable loci, ITS1, ITS2, and 28S, were aligned with the E-INS-i strategy using ‘--genafpair --maxiterate 1000’. In parallel, we generated structure-guided alignments for each locus using the MAFFT X-INS-i framework (mafft-xinsi) with default structural-alignment settings^32^. Poorly aligned and highly entropic columns were removed with BMGE ^52^ using the following parameters: ‘-g 0.7 -h 0.6’ for ITS sequences and ‘-g 0.5 -h 0.5’ for rRNA sequences. For phylogenetic inference, maximum-likelihood trees were reconstructed with IQ-TREE 3^53^ separately for each locus and alignment strategy. We used the built-in ModelFinder procedure (‘-m MFP’) to select an appropriate nucleotide substitution model and IQ-TREE default optimization settings; branch support was estimated via IQ-TREE’s default bootstrap scheme (1,000 ultrafast bootstrap replicates and 1,000 SH-aLRT replicates; ‘-bb 1000 -alrt 1000 -bnni’). The resulting best-tree files were used both as final phylogenies and as inputs to comparative tree analyses.

### Tree-level clade clustering assessment

To compare broad phylogenetic organization across loci and tree-building strategies, we evaluated each inferred tree against the expected vertebrate clade assignments of its terminal taxa rather than against a single external reference topology. Before analysis, trees were restricted to the overlap species set, and each terminal taxon was annotated with its clade identity. Clade labels were then projected onto the terminal order of the plotted rectangular tree and used to calculate two descriptive clustering statistics. First, the ‘adjacent same-group fraction’ was defined as the proportion of adjacent terminal-taxon pairs sharing the same clade label; higher values indicate stronger local clustering of taxa from the same clade. Second, the ‘run fragmentation index’ quantified the extent to which same-clade blocks were broken up along the terminal order. Contiguous runs of identical clade labels were counted across the ordered tips, and the excess number of runs above the theoretical minimum, one run per represented clade, was normalized by the maximum possible excess runs for the observed number of tips. This index ranges from 0 to 1, with values near 0 indicating stronger clade-level contiguity and values near 1 indicating greater intermixing among clades. These metrics provided a benchmark-like summary of how well each tree recapitulated broad expected clade structure, while avoiding reliance on a fully specified reference topology.

### Tree comparison and quartet distances

To quantify topological similarity between trees derived from different units (ITS1, ITS2, and 45S rRNA), we computed normalized quartet distances for all tree pairs on the intersection of shared taxa using R package “Quartet”^54^. For each tree pair we calculated the normalized quartet distance (number of disagreeing quartets divided by the total number of quartets) using established quartet-distance implementations. We constructed co-phylogenetic “tanglegram” plots using the ‘cophylo’ function from the “phytools” R package^55^, matching species between trees and drawing corresponding tips as linked pairs.

### Sequence–structure divergence analysis

Representative consensus sequences were aligned using MAFFT v7.526^51^ with automatic strategy selection. Pairwise nucleotide divergence was estimated from the alignment using the Kimura two-parameter model implemented in EMBOSS distmat. Secondary structures for each ITS2 sequence were predicted using RNAfold from the ViennaRNA package^56^, and pairwise structural distances were computed using RNAdistance -Xm, which reports base-pair distance between folded structures. Correlations between pairwise sequence divergence and structural divergence were calculated across all taxon pairs. Because absolute base-pair distance scales in part with RNA length, structural distances were also normalized by the mean sequence length of each species pair to assess sequence–structure coupling independently of spacer size.

### Statistical environment and visualization

All data processing and statistics were performed in R (version 4.5.2) using base R and the tidyverse^57^ for data manipulation and ggplot2^58^ for plotting. Where indicated, quasi-random jittering of points over violins used ggbeeswarm ^59^ to improve visibility in dense regions. Custom R and Python scripts were used to parse GFF files, compute rDNA unit sizes, summarize per-species statistics, and generate figures.

## Notes

https://www.dropbox.com/scl/fi/3payv9ncaidba3h1iiz1u/SupplementaryTables.xlsx?rlkey=44852pkqdtdidbb08qsu3hlag&dl=0

### Competing Interest Statement

The authors have declared no competing interest.

### Summary of Updates

Some references (including the VGP flagship manuscript, Formenti et al. 2026) required updating. Also, the previous version with figures embedded in the text produced a PDF with poor quality of the illustrations. The revision has figures uploaded separately to (hopefully) increase the quality.

